# Protein model accuracy estimation empowered by deep learning and inter-residue distance prediction in CASP14

**DOI:** 10.1101/2021.01.31.428975

**Authors:** Xiao Chen, Jian Liu, Zhiye Guo, Tianqi Wu, Jie Hou, Jianlin Cheng

## Abstract

The inter-residue contact prediction and deep learning showed the promise to improve the estimation of protein model accuracy (EMA) in the 13th Critical Assessment of Protein Structure Prediction (CASP13). During the 2020 CASP14 experiment, we developed and tested several EMA predictors that used deep learning with the new features based on inter-residue distance/contact predictions as well as the existing model quality features. The average global distance test (GDT-TS) score loss of ranking CASP14 structural models by three multi-model MULTICOM EMA predictors (MULTICOM-CONSTRUCT, MULTICOM-AI, and MULTICOM-CLUSTER) is 0.073, 0.079, and 0.081, respectively, which are ranked first, second, and third places out of 68 CASP14 EMA predictors. The single-model EMA predictor (MULTICOM-DEEP) is ranked 10th place among all the single-model EMA methods in terms of GDT_TS score loss. The results show that deep learning and contact/distance predictions are useful in ranking and selecting protein structural models.

## 1. Introduction

In a protein structure prediction process, the estimation of model accuracy (EMA) or model quality assessment (QA) is important for selecting good tertiary structure models from many predicted models. EMA also provides valuable information for users to apply protein structural models in biomedical research. In the 13th Critical Assessment of Protein Structure Prediction (CASP13), the inter-residue contact information and deep learning were the key for DeepRank (Hou et al. 2019) to achieve the best performance in ranking protein structural models with the minimum loss of GDT-TS score (Zemla 2003). Recently, inter-residue distance predictions have been used with deep learning for the estimation of model accuracy (Shuvo, Bhattacharya, and Bhattacharya 2020, Jing and Xu 2020, Hiranuma et al. 2020).

To further investigate how residue-residue distance/contact features may improve protein model quality assessment, we developed several EMA predictors based on different ways of using contact and distance predictions as features with deep learning and tested them in the 2020 CASP14 experiment. Some of these predictors are based on the features used in our CASP13 EMA predictors, while others use new contact/distance-based features (Chen et al. 2020) or new image similarity-derived features by treating predicted inter-residue distance maps as images and calculating the similarity between the distance maps predicted from protein sequences and the distance maps directly computed from the 3D coordinates of a model. All the methods predict a GDT-TS score normalized between 0 and 1 for a model of a target, which measures the quality of the model.

According to the nomenclature in the field, these CASP14 MULTICOM EMA predictors can be classified into two categories: *multi-model methods* (MULTICOM-CLUSTER, MULTICOM-CONSTRUCT, MULTICOM-AI, MULTICOM-HYBRID) that use some features based on the comparison between multiple models of the same protein target as input and *single-model methods* (MULTICOM-DEEP and MULTICOM-DIST) that only use the features derived from a single model without referring to any other model of the target. Multi-model methods had performed better than single-model methods in most cases in the past CASP experiments. However, multi-model methods may perform poorly when there are only a few good models in the model pool of a target, while the prediction of single-model methods for a model is not affected by other models in the pool. Moreover, single-model methods can predict the absolute quality score for a single protein model (Wang et al., 2009, Cao et al., 2014), while the score predicted by multi-model methods for a model depends on other models in the model pool. In the following sections, we describe the technical details of these two kinds of methods, analyze their performance in the CASP14 experiment, and report our findings.

## 2. Method

### 2.1. The pipeline for estimation of model accuracy

When a protein target sequence and a pool of predicted structural models for the target are received, a MULTICOM EMA predictor call an inter-residue distance predictor - DeepDist (Wu et al. 2020) and/or an inter-residue contact predictor - DNCON2 (Adhikari et al., 2018) / DNCON4 (Wu et al., 2019) - to predict the distance map and/or contact map for the target. Some distance/contact-based features such as the Pearson’s correlation between the predicted distance map (PDM) and a structural model’s distance map (MDM), the percentage of predicted inter-residue contacts (i.e., short-range, medium-range, and long-range contacts) occurring in the structural model are generated. Several metrics of measuring the similarity between images including the distance-based DIST descriptor (Oliva and Torralba 2001), Oriented FAST and Rotated BRIEF (ORB) (Rublee et al. 2011), and PHASH (Karasikov et al. 2019), and PSNR & SSIM (Hore and Ziou 2010) as well as root mean square error (RMSE) are used to calculate the similarities or difference between PDM and MDM as features.

Other non-distance/contact features used in DeepRank are also generated for the predictors, which include single-model features, i.e., SBROD (Karasikov et al. 2019), OPUS_PSP (Lu, Dousis, and Ma 2008), RF_CB_SRS_OD (Rykunov and Fiser 2007), Rwplus (Zhang and Zhang 2010), DeepQA (Cao et al. 2016), ProQ2 (Uziela and Wallner 2016), ProQ3 (Uziela et al. 2016), Dope (Shen and Sali 2006), Voronota (Olechnovič and Venclovas 2014), ModelEvaluator (Wang et al., 2008), QMEAN (Benkert et al., 2011), solvent accessibility score (i.e., solvent) generated by SSpro4 (Magnan & Baldi, 2014) and DSSP (Kabsch et al. 1983), regular secondary structure (helix and beta sheet) penalty score (i.e, ss_penalty), secondary structure similarity score (i.e, ss_sim) (Wang et al., 2008), paired euclidean distance score (i.e, euclidean), total surface score (i.e., total_surface), weighted exposed surface area score (i.e., weighted_exposed), and an average feature probability density score (i.e. feature_probability) (Cao, and Cheng 2016). The three multi-model features are APOLLO (Wang, Eickholt, and Cheng 2011), Pcons (Wallner and Elofsson 2006), and ModFOLDClust2 (McGuffin and Roche 2010). Different combinations of the features described above are used with deep learning to predict the GDT-TS score of a model, resulting in multiple MULTICOM EMA predators. **Table 1** is the summary of the features employed by them.

**Table 1:**
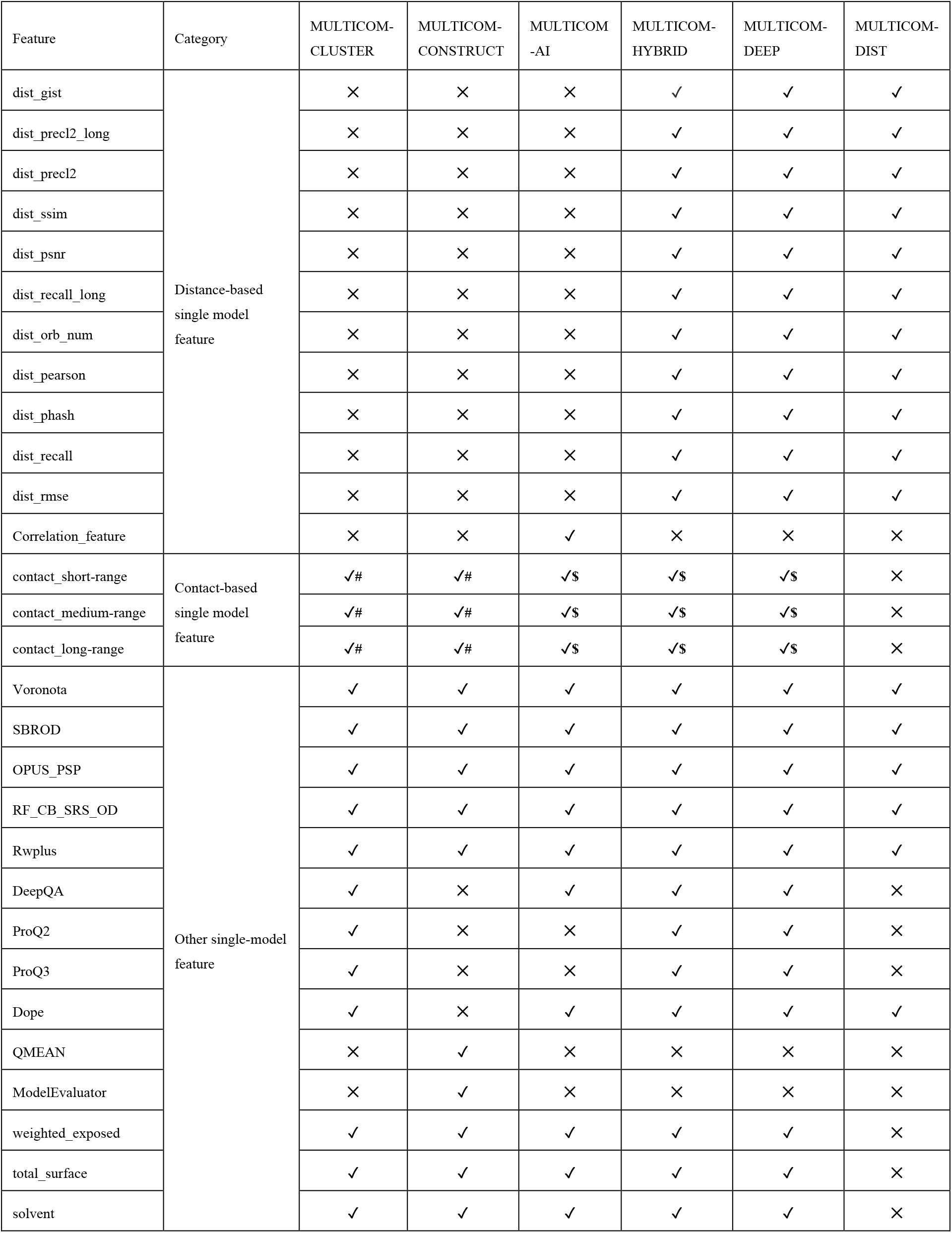
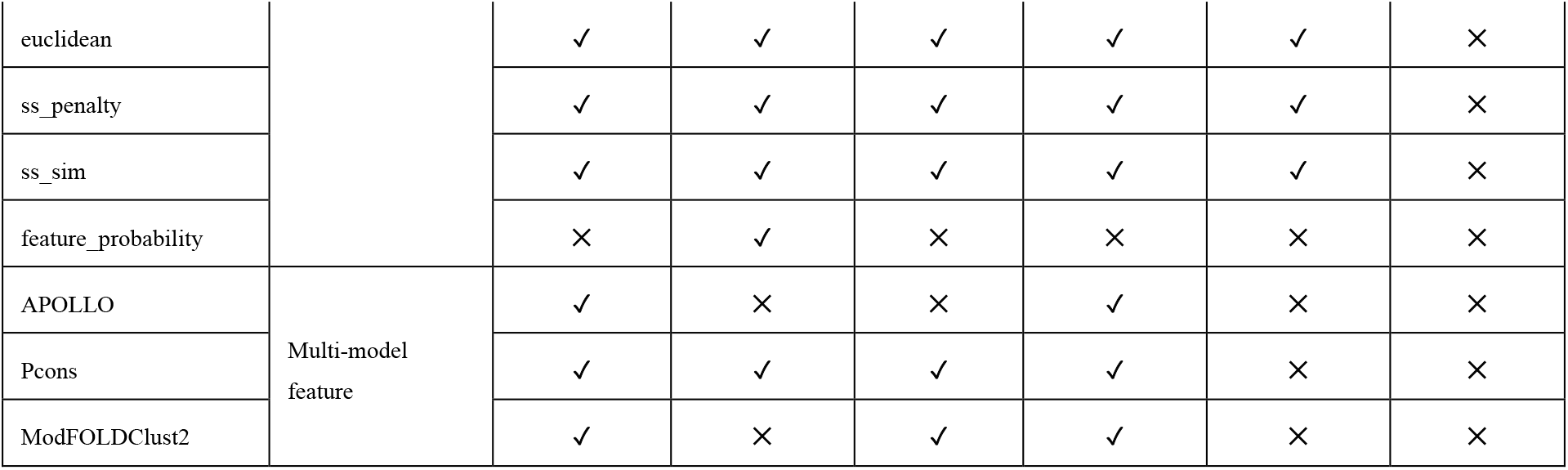
Features used by six MULTICOM EMA predictors. The features are divided into four categories: distance-based single-model features, contact-based single-model features, other single-model features, and multi-model features. ✓: a feature used by a predictor. ✕ a feature not used by a predictor. #: features based on contacts predicted by DNCON2. $: features based on contacts predicted by DNCON4.

In order to handle a partial structural model that contains only the coordinates for a portion of the residues of a target, the ratio of the residues in the model and the total number of the residue of the target (i.e., target sequence length) is calculated. The predicted score for a partial model by a deep learning predictor times the ratio is its final predicted quality score of the partial model.

Another caveat is that, because generating some features such as pairwise similarity features between models for very large protein targets (i.e., T1061, T1080, T1091) is very time consuming, the MULTICOM predictors use the domain prediction to divide their models into multiple domains first. Then the pipeline described above is used to predict the quality score of the models of each domain. The average score of multiple domains of a full-length structural model is used as its estimated model quality score.

### 2.2. Deep learning training and prediction

The structural models of CASP8-CASP12 were used as a training dataset to train and validate the deep learning EMA methods. The structural models of CASP13 were used as a test dataset to evaluate the methods before they were blindly tested in CASP14.

The training dataset was split into K equal-size folds. Each fold was used as the validation set for parameter tuning while the remaining K-1 folds were used to train a deep learning model to predict the quality score. This process was repeated K times, yielding K trained EMA predictors each validated on one-fold (i.e., K-fold cross-validation) (Hou et al. 2019). The average output of the K predictor for a model can be used as the final predicted quality score of the model (i.e., the ensemble prediction).

Moreover, some MULTICOM EMA predictors used a two-stage training strategy, adding another round of training (stage-2 training) on top of the modeling trained above (i.e., stage-1 training). The original input features of a model were combined with the K quality scores of the model predicted by the K EMA predictors from the stage-1 training as input features for another deep learning model to predict its final quality score. The deep learning model was trained on the same set of models of CASP8-12 in the stage-2 training.

### 2.3. Implementation of MULTICOM EMA predictors

MULTICOM-CONSTRUCT and MULTICOM-CLUSTER use the same deep learning architecture and input features as in DeepRank (Hou et al. 2019). They were trained the same 10-fold cross-validation. However, MULTICOM-CONSTRUCT uses the average output of the 10 predators trained in the stage-1 training as prediction, but MULTICOM-CLUSTER applies the deep learning model trained with the two-stage training strategy to make prediction. MULTICOM-AI is built upon a variant of DeepRank (Chen et al. 2020). Its deep network was trained with 5-fold cross-validation.

MULTICOM-HYBRID, MULTICOM-DEEP, and MULTICOM-DIST use some input features that are quite different from MULTICOM-CONSTRUCT, MULTICOM-CLUSTER and MULTICOM-AI (see **Table 1**). First, DNCON2 is replaced with DNCON4 to make contact predictions for contact-based features. The inter-residue distance-based features (i.e., SSIM & PSNR, GIST, RMSE, Recall, Precision, PHASH, Pearson correlation, ORB) calculated from distance maps predicted by DeepDist are also used as their input features. MULTICOM-HYBRID uses the new contact-based features and the new distance-based features as well as the nine single-model quality scores and the three multi-model features of DeepRank to make predictions. Because it uses the multi-model features as input, it is a multi-model method. MULTICOM-DEEP uses the same features as MULTICOM-HYBRID except that the three multi-model features are removed. Therefore, MULTICOM-DEEP is a single-model method. MULTICOM-DIST is a light version of MULTICOM-DEEP. It uses a subset of the features of MULTICOM-DEEP, excluding several features including DeepQA, ProQ2, ProQ3, contact matching scores that take quite some time to generate. MULTICOM-HYBRID, MULTICOM-DEEP, MULTICOM-DIST were trained on CASP8-12 protein models and tested on CASP13 models, yielding the similar GDT-TS loss on the test dataset (i.e., 0.0487, 0.0533, 0.0503, respectively).

## 3. Result and Discussion

### Evaluation data and metrics

MULTICOM EMA predictors blindly participated in the CASP14 EMA prediction category from May to July 2020. CASP14 evaluated EMA methods in two stages (Cheng et al. 2019). In stage 1, 20 models of each target with very different quality were sent to the registered EMA predators to predict their quality. In stage 2, top 150 models selected by a simple consensus EMA method for each target were used for the EMA predictors to predict their quality. Because CASP14 only released the evaluation results on stage-2 models, we analyze our methods on the stage-2 models in this study. CASP4 used 69 valid targets to evaluate EMA predictors. Each target may have one domain or more. The domains are classified into three categories: (1) *template-based modeling (TBM) domains* - the regular domains that have known structure templates in the Protein Data Bank (PDB) (Berman, H. M. et al. 2000) (TBM domains are further classified into TBM-easy and TBM-hard categories according to the difficulty of predicting their tertiary structures); (2) *free modeling (FM) domains* - the very hard domains that do not have any known structure templates in the PDB; and (3) *something between the two (FM/TBM)*, which may have some very weak templates that cannot be recognized by existing template-identification methods. If a target contains multiple domains of different difficulty categories, it is classified into the most difficult category of its domains in this study.

We downloaded the official evaluation results of the CASP14 EMA predictors from the CASP14 website, analyzed MULTICOM EMA predictors’ performance, and compared them with other EMA predictors. We use the average loss of GDT-TS score of an EMA predictor over all the CASP14 targets as the main metric to evaluate its performance. The GDT-TS loss of a predictor on a target is the absolute difference between the true GDT-TS score of the No. 1 model selected from all the models of the target by the predicted GDT-TS scores and the true GDT-TS score of the best model of the target. A loss 0 means the best model in the model pool of a target is chosen by an EMA predictor, which is the ideal situation. The average GDT-TS loss of a predictor on all the CAS14 targets is used to evaluate how well it can select or rank protein models. In addition to the GDT-TS loss, we also use the average Pearson’s correlation between the predicted GDT-TS scores of the models of a target and their true GDT-TS scores over the CASP14 targets to evaluate the EMA methods.

#### 3.1. GDT-TS loss and Pearson’s correlation of the MULTICOM EMA predictors in CASP14

Boxplots in **Figure 1(A)** show the GDT-TS loss of each target, average loss, and variation of the loss for the six MULTICOM EMA predictors on all the CASP14 targets. MULTICOM-CONSTRUCT, MULTICOM-AI, MULTICOM_CLUSTER, MULTICOM-HYBRID, MULTICOM-DEEP and MULTICOM-DIST attain 0.0734, 0.0792, 0.0806, 0.086, 0.104, and 0.124 (GDT_TS) loss on average, respectively. Overall, the four multi-model methods (MULTICOM-CONSTRUCT, MULTICOM-AI, MULTICOM_CLUSTER, MULTICOM-HYBRID) perform better than the two single-model methods (MULTICOM-DEEP and MULTICOM-DIST), among which MULTICOM-CONSTRUCT perform best in terms of the GDT-TS loss.

**Figure 1.**
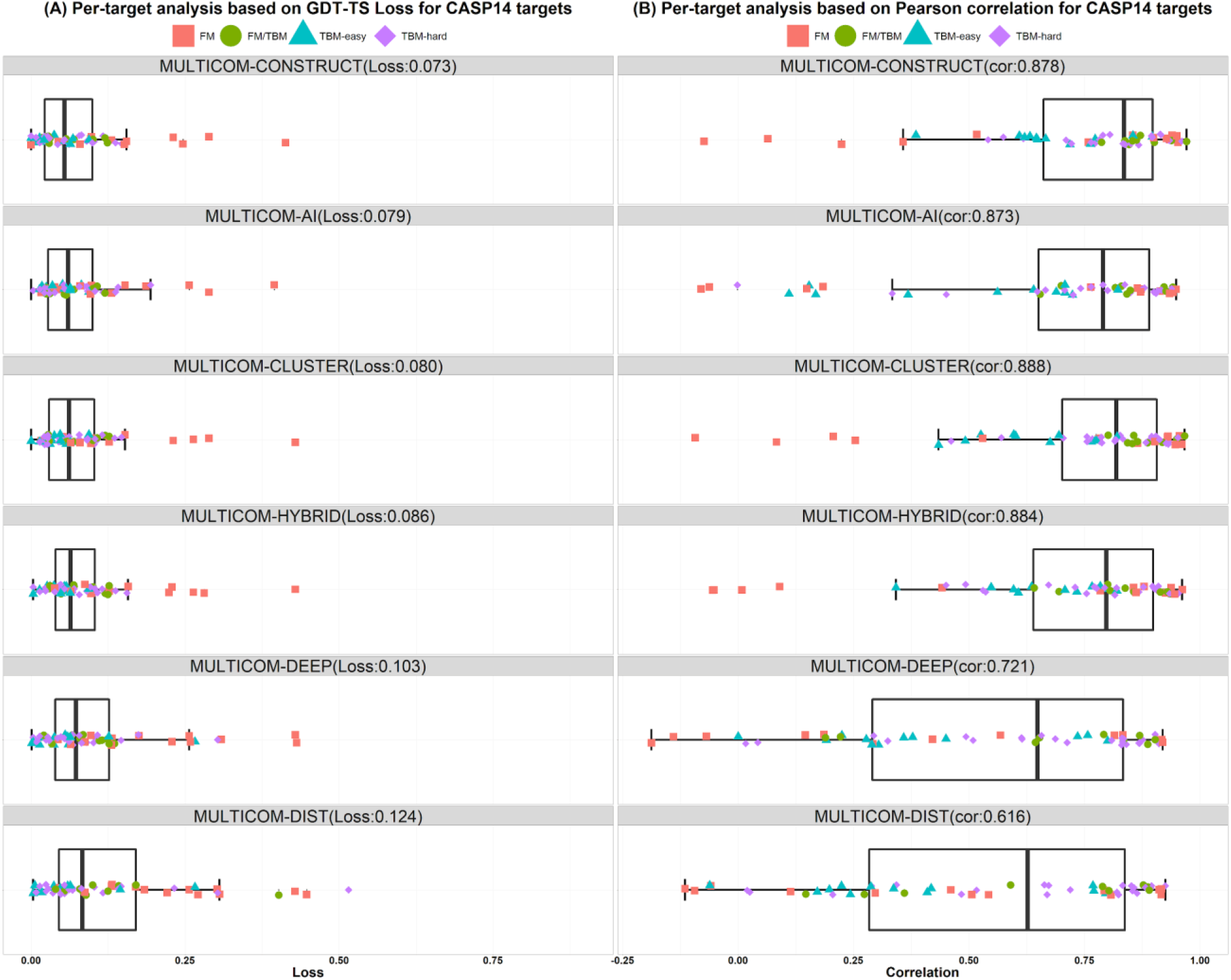
The boxplots of MULTICOM predictors’ performance on CASP14 targets. (A) GDT_TS score loss. (B) Pearson’s correlation score. Different colors/shapes denote different kinds of targets.

**Figure 1(B)** plots the Pearson’s correlation scores of the CASP14 targets for each MULTICOM predictor. MULTICOM-CLUSTER obtains the highest global correlation coefficient (0.888) and MULTICOM-DIST’s correlation coefficient (0.616) is the lowest. The four multi-model methods’ correlation scores are close (i.e., MULTICOM-CONSTRUCT: 0.878, MULTICOM-AI: 0.873, MULTICOM-HYBRID: 0.884, MULTICOM-CLUSTER: 0.888). Their high correlation scores indicate a strong positive correlation relationship between the true GDT_TS scores and predicted GDT_TS scores. The two single-model methods perform worse than the multi-model methods. MULTICOM-DEEP, MULTICOM-DIST achieve a correlation score of 0.721 and 0.611, respectively.

#### 3.2. Comparison between multi-model methods and single-model methods

**Figure 2(A)** illustrates six MULTICOM EMA predictor’s performance in each target category. The multi-model methods consistently outperform the single-model methods in all the categories, indicating that there is still a significant room for single-model methods to improve. For instance, on the TBM-easy targets, four multi-model methods have very close GDT_TS loss (~0.04), which is 33.3% lower than the single-model methods’ loss (~0.06). MULTICOM-CONSTRUCT obtains the lowest loss (0.057) on TBM-hard targets, while MULTICOM-DIST gets the highest loss (0.103). On the FM/TBM targets, MULTICOM-CONSTRUCT has the lowest loss of 0.058, 34% lower than MULTICOM-DEEP’s 0.09. On the most challenging FM targets, MULTICOM-AI has the lowest loss of 0.145, while MULTICOM-DEEP and MULTICOM-DIST get 0.203 and 0.229 loss respectively. The results show the GDT-TS loss is lower on easier targets than harder targets for all the MULTICOM EMA predictors generally, indicating that it is still easier to rank the models of easy targets than hard targets.

**Figure 2.**
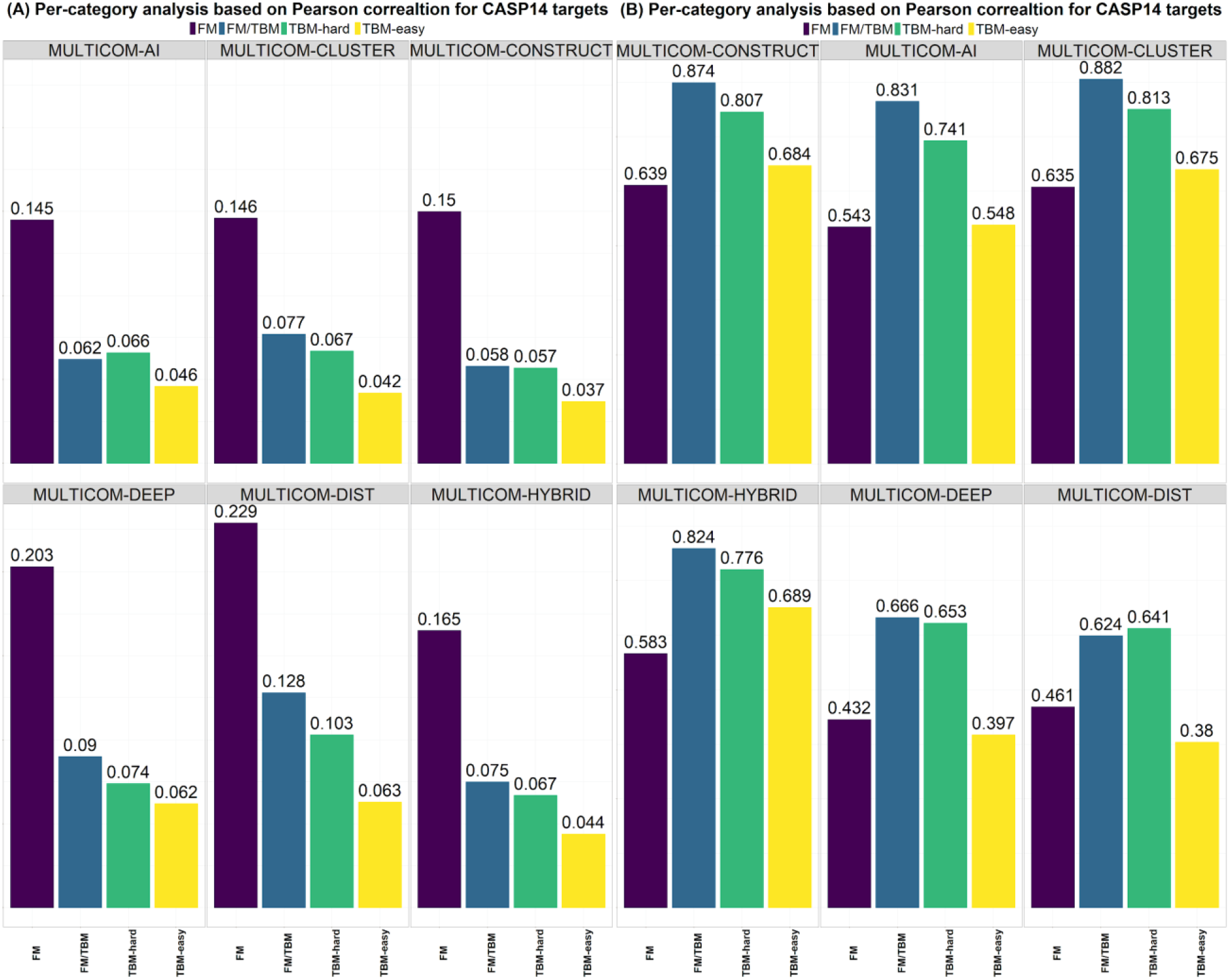
The performance of MULTICOM EMA predictors on four different categories of targets (FM, FM/TBM, TBM-hard, and TBM-easy). (A) GDT_TS ranking loss. (B) Pearson’s correlation.

**Figure 2(B)** shows the largely similar trend in Pearson’s correlation evaluation. The four multi-model methods perform better than the two single-model methods. On FM targets, MULTICOM-CONSTRUCT achieves the highest correlation coefficient (0.639) and MULTICOM-CLUSTER gets a very similar correlation score (0.635). MULTICOM-CONSTRUCT’s correlation score is 39% higher than that of MULTICOM-DEEP (0.432). On FM/TMB targets, MULTCOM-CLUSTER result is the best (0.874), 40% higher than MULTICOM-DIST’s score (0.624). MULTICOM-CLUSTER attains 0.813 correlation coefficient on TBM-hard targets and MULTICOM-DIST has lowest correlation coefficient (0.641). On the easiest TBM-easy targets, MULTICOM_CONSTRUCT, MULTICOM-CLUSTER and MULTICOM-HYBRID have the correlation score of around 0.68, while two single-model methods perform worse on these targets. But there is some difference between the evaluation based on the correlation and the GDT_TS ranking loss. The predictors achieve the best performance on FM/TBM targets according to the correlation, but not on the easiest TBM-easy targets according to the ranking loss.

#### 3.4 Comparison with Other CASP14 EMA Predictors

CASP14 released the whole server’s overall and per-target performance in ranking structural models. **Table 2** is the summary of the GDT_TS loss and the Local Distance Difference Test (LDDT) (Mariani et al., 2013) loss of top 20 out of 68 predictors that predicted the quality of the models of most CASP14 targets. MULTICOM-CONSTRUCT, MULTICOM-AI, MULTICOM-CLUSTER, MULTICOM-HYBRID is ranked first, second, third, and tenth in terms of GDT-TS loss, respectively. MULTICOM-DEEP was at 20th among all the EMA predictors and 8th among the single-model EMA predictors (data not shown). Different from GDT-TS loss, in terms of LDDT loss, the ranks of MULTICOM predictors are lower. Four MULTICOM predictors are ranked among top 20, and MULTICOM-CONSTRUCT is ranked sixth. The lower ranking of MULTICOM EMA predictors in terms of LDDT loss is partially because they were trained to predict GDT-TS score instead of LDDT score.

**Table2.**
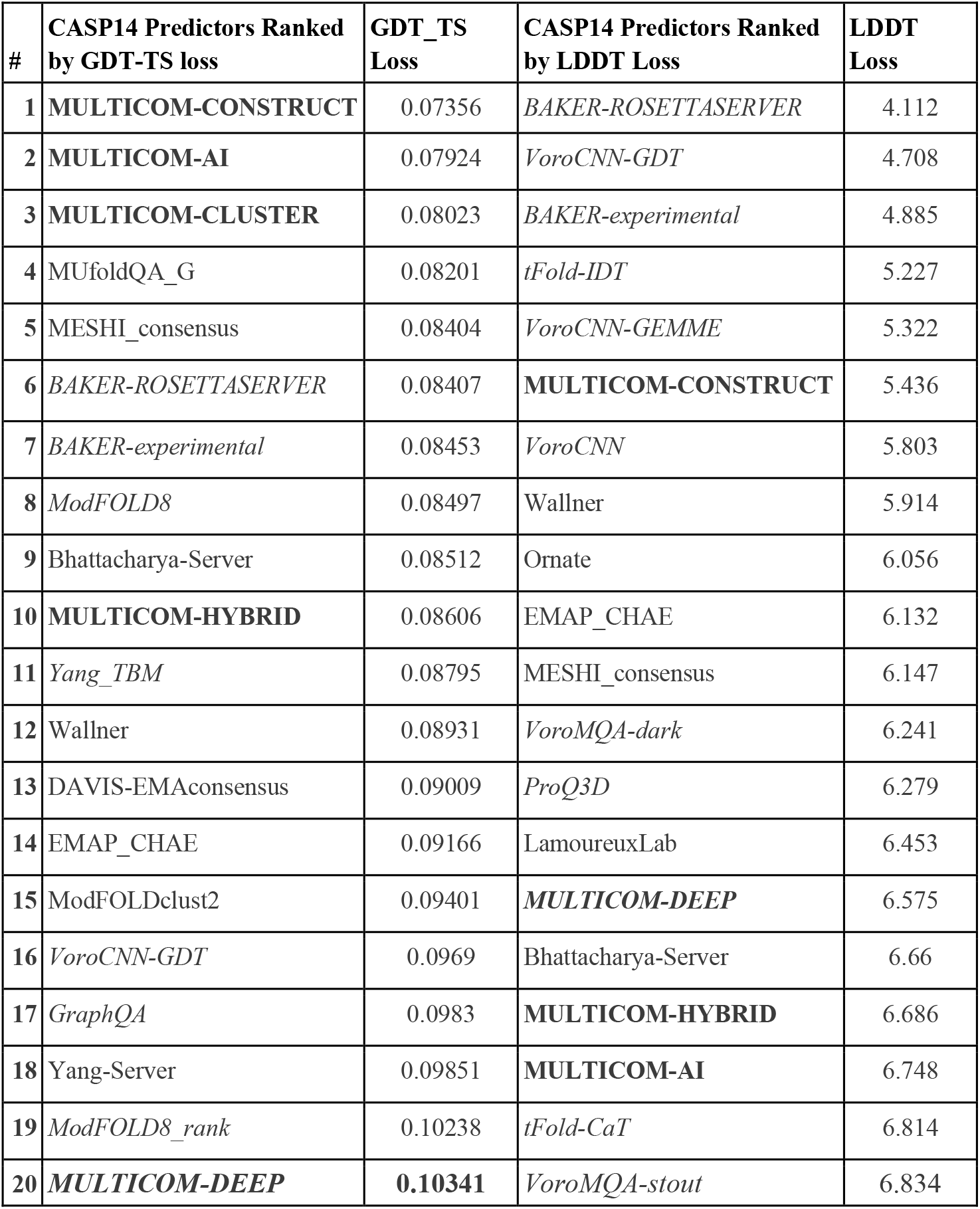
Top 20 CASP10 EMA Predictors ranked by GDT-TS loss and LDDT Loss, respectively. Here, the original, official GDT-TS loss of each predictor is normalized into the range [0, 1]. Bold font stands for MULTICOM predictors. Italic font denotes single-model methods.

#### 3.5 Case Study

One successful prediction example made by MULTICOM EMA predictors is shown in **Figure 3**. The predicted distance map is similar to the true distance map of the target. Three MULTICOM EMA predictors (i.e., MULTICOM-AI, MULTICOM-CONSTRUCT, MULTICOM-CLUSTER) successfully rank the best model at the top, resulting in a loss of 0.

**Figure 3.**
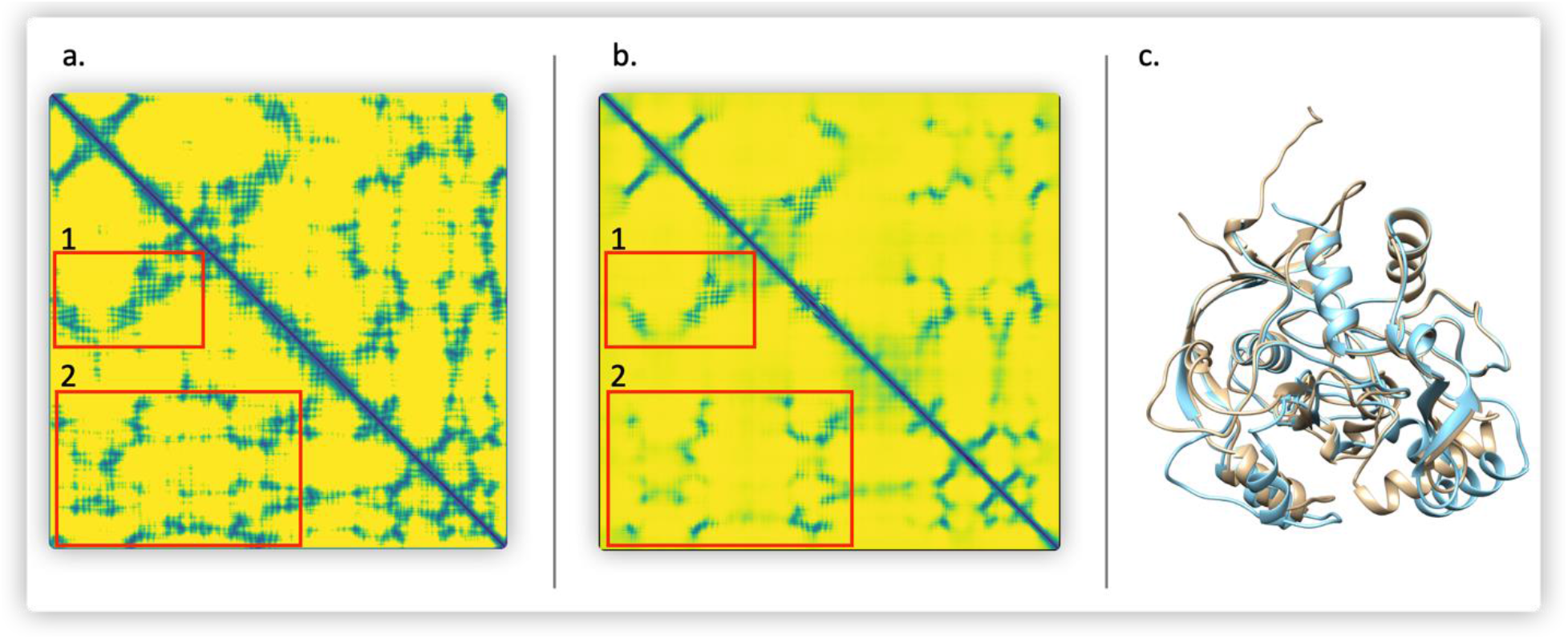
A good EMA example (T1028). (A) true distance map (darker collar means shorter distances). (B) predicted distance map. (C) top model selected by a MULTICOM predictor (MULTICOM-AI) (light blue) versus true structure (light yellow). The GDT-TS ranking loss is 0. Red rectangles in the maps highlight some long-range contacts.

**Figure 4** illustrates a failed example (T1039), on which MULTICOM-AI has a high GDT-TS ranking loss of 0.289. The precision of top L/2 long-range contacts (L: sequence length) calculated from the predicted distance map is only 18.52%. The predicted distance map and the true distance map of this target are very different. Particularly, a lot of long-range contacts are not predicted.

**Figure 4.**
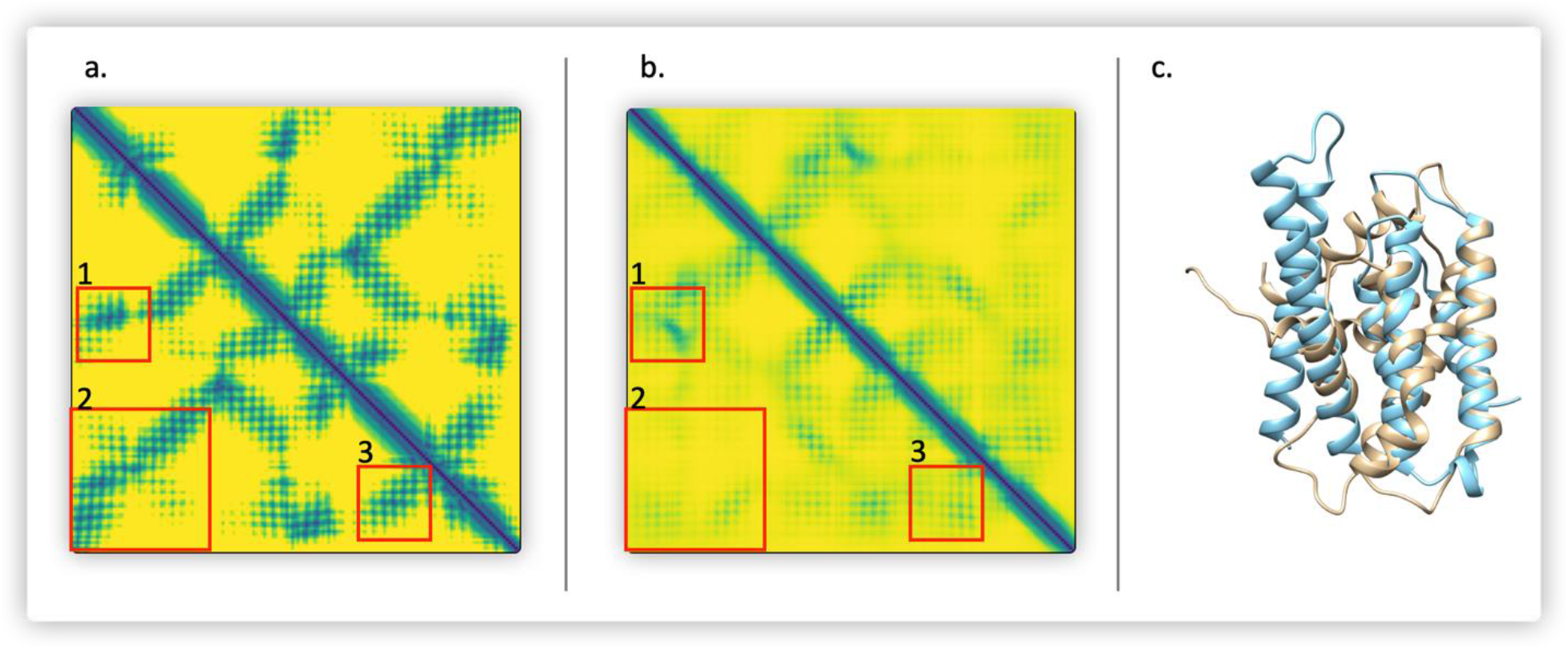
A failed example (T1039). (A) the true distance map. (B) the predicted distance map. (C) top model selected by MULTICOM-AI (light blue) versus true structure (light yellow). Red boxes highlight regions that the true map and the predicted map differ a lot. The GDT-TS loss of MULTICOM-AI is 0.289.

#### 3.6 Discussion

The average GDT-TS loss of our best MULTICOM EMA predictors on the CASP14 dataset (e.g., ~0.07 −0.08) is higher than the average loss (e.g., ~0.5) on the CASP13 dataset, suggesting that it is harder to rank the CASP14 models than the CASP13 models. It may be due to the fact most models of CASP13 or even CASP8-12 were generated by the traditional modeling methods, but most models in CASP14 were generated by new distance-based protein structure modeling methods such as trRosetta developed after CASP13, which may have some different properties from the CASP8-13 models. Therefore, the generalized prediction performance of MULTICOM EMA predictors trained on CASP8-12 models performed worse on CASP14 models than CASP13 models. Therefore, it is important to create larger model datasets created by the new model generation methods to train EMA methods in the future. This may also partially explain why the new distance-based features used in MULTICOM-HYBRID, MULTICOM-DEEP, and MULTICOM-DIST do not seem to improve the performance of EMA over the MULTICOM predictors using only contact-based features on the CASP14 dataset, even though they are mostly accurate enough to build the tertiary structural models according to our tertiary structure prediction experiment in CASP14. Another reason is that, because the distance prediction had been used to generate some tertiary structure models included in the CASP14 models, using the features based on the same distance information in the model accuracy estimation again might create an artificial bias to rank those models higher, leading to the higher loss. Therefore, using the distance information predicted by a distance predictor in model accuracy estimation different from the distance predator used in tertiary structure prediction is desirable. Moreover, further improving the accuracy of distance predictions, particularly for hard FM targets, is useful to improve their effects on EMA. Finally, instead of using the expert-curated features derived from distance maps as input, directly using raw distance maps or even raw 3D structural models themselves as input may allow deep learning to automatically figure out the relevant features in the input to estimate model accuracy more effectively.

## 4. Conclusion and Future Work

We developed several deep learning EMA predictors, blindly tested them in CASP14, and analyzed their performance. Our multi-model EMA predictors performed best in CASP14 in terms of the average GDT-TS loss of ranking protein models. The single-model EMA predators using inter-residue distance features also delivered a reasonable performance on most targets, indicating the distance information is useful for protein model quality assessment. However, estimating the accuracy of models of some hard targets remains challenging for all the methods. The better ways of using distance features, more accurate distance prediction for hard targets, and larger training datasets generated by the latest protein tertiary structure prediction methods in the field are needed to further improve the performance of estimating model accuracy. Moreover, instead of predicting one kind of quality score (e.g., GDT-TS or LDDT score) for a structural model, it is desirable to predict multiple quality scores via multi-task machine learning to meet the different needs in different situations.

## Acknowledgements

The project is partially supported by two NSF grants (DBI 1759934 and IIS1763246), one NIH grant (GM093123), two DOE grants (DE-SC0020400 and DE-SC0021303), and the computing allocation on the Summit supercomputer provided by Oak Ridge Leadership Computing Facility (Project ID: BIF132).

